# Multiphoton microscopy for label-free multicolor imaging of peripheral nerve

**DOI:** 10.1101/2021.09.30.462641

**Authors:** Lars Rishøj, Iván Coto Hernández, Siddharth Ramachandran, Nate Jowett

## Abstract

Conventional histomorphometry of peripheral nerve entails lengthy chemical processing, ultrathin sectioning in resin, and imaging by light or electron microscopy. Multiphoton microscopy techniques exist enabling label-free and in vivo imaging of histological samples. Third-harmonic-generation microscopy has recently been demonstrated effective for imaging the myelin sheath of peripheral nerve axons in animal models. Herein, we characterize use of second and third harmonic generation microscopy for label-free imaging of murine and human peripheral nerve via a novel multicolor multiphoton microscope based on a single excitation wavelength at 1300 nm. Second harmonic generation signal from collagen centered about 650 nm delineates neural connective tissue, while third harmonic general signal centered about 433 nm delineates myelin and other lipids. In transgenic mice expressing yellow fluorescent protein linked to the *thy1* promoter, three-photon-excitation with emission peak at 527 nm delineates axoplasm. We compare label-free multiphoton imaging of murine and human peripheral nerve against conventional chemical stains and discuss clinical implications of this approach in guiding intraoperative decision making in nerve transfer procedures.

## Introduction

Peripheral nerve injury may result in devastating functional and aesthetic sequelae. Nerve transfer procedures may be employed for rehabilitation of peripheral nerve deficits. These surgical procedures comprise re-routing axons from less important neighboring nerves towards critical distal targets. During an end-to-end nerve transfer procedure, a donor nerve is dissected, transected distally, and proximal stump transposed to meet the distal stump of the recipient nerve without tension. Examples of nerve transfer procedures include transfer of sensory nerves supplying the forehead or earlobe to the cornea in the management of neurotrophic keratitis^1–3^, and transfer of ipsilateral trigeminal nerve motor branches or select contralateral facial nerve branches in the management of unilateral facial palsy^4^.

Donor nerve myelinated axonal load is a critical factor in functional outcomes following motor nerve transfer procedures^5,6^. In cross-facial nerve transfer for smile reanimation, a minimum of 900 myelinated axons in the donor nerve branch is thought necessary to reduce the risk of procedural failure^7^. Presently, intraoperative donor nerve selection is based on surgeon experience and rough estimation of fiber counts based on nerve branch calibre^8^. Though selection of larger caliber donor nerves increases the likelihood of sufficient target muscle neurotization, it must be weighed against the risk of weakening the target musculature of the donor nerve. A means for rapid and reliable quantification of axons present within donor nerve branches could inform operative decision making. As end-to-end nerve transfer procedures such as cross-facial nerve grafting require sectioning of the donor nerve, opportunity exists for intraoperative cross-sectional histopathologic analysis to calculate donor nerve myelinated axon counts. Where the number of myelinated axons present in a given donor branch is suboptimal, an additional donor branch may be selected to achieve adequate neurotization of the distant target.

Conventional nerve histomorphometry entails lengthy sectioning and chemical staining steps, rendering it impractical for intraoperative use. We recently characterized means for quantification of myelinated axon counts from frozen sections of peripheral nerve using a non-toxic fluorescent dye^9^. Several label-free nonlinear optical microscopy (NLOM) approaches for imaging of peripheral nerve myelin sheaths have also been described, including stimulated Raman scattering (SRS)^10^ paired with coherent anti-Stokes Raman scattering (CARS) microscopy, and third harmonic generation (THG) microscopy^11^.

NLOM is an alternative to conventional confocal microscopy for the study of the three-dimensional samples^12,13^, comprising multiphoton fluorescence and harmonic generation techniques. Multiphoton excitation fluorescence microcopy is based on the absorption of two or more long wavelength photons to excite a molecule, with subsequent decay to a lower energy state via spontaneous emission of a single shorter wavelength photon. Harmonic generation microscopy is a non-absorptive optical process wherein incident photons from a high intensity laser pulse are converted by interaction with a non-linear material into harmonic frequencies having integer multiples of the incident wavelength. The evolution of harmonic generation microscopy mirrors that of multiphoton excitation microscopy as the techniques employ similar microscope hardware. Second harmonic generation (SHG) and THG microscopy techniques enable label-free high-contrast imaging of specific constituents of biological tissue with minimal or absent need for sample preparation. SHG microscopy can image fibrillar collagen, and is typically performed using excitation wavelengths between 700-1000 nm. THG microscopy can image lipids, and may be achieved using extended range excitation from a Ti:Sapphire laser coupled to an optical parameter oscillator (OPO) in the range of 1050-1250 nm^14^. Recently, improved penetration depths with label-free THG microscopy have been achieved using excitation wavelengths in the second optical window of biological tissue between 1050 nm and 1350 nm^15^.

Recently, we developed a novel high peak power (1.1 MW, 80 nJ and 74 fs) fiber laser at 1300 nm with selectable repetition rate between 1-10 MHz^16^. The operational wavelength of this novel fiber laser is optimal for three-photon microscopy of biological tissue, for which no commercial microscope currently exists. Using this fiber laser as an illumination source, we herein developed a microscope optimized for multi-color NLOM. The custom design of this microscope enables maximal efficiency of two and three-photon NLOM^17,18^. We employed the microscope for label-free multicolor imaging of human and murine peripheral nerve ^19^. Compared to conventional nerve histology, harmonic imaging provides high specificity while avoiding the need for chemical staining and its associated toxicity. Collagen-dense nerve sheaths (endoneurium, perineurium, and epineurium) are resolved using SHG signal centered at ~650 nm, while lipid-dense myelin sheaths of individual myelinated axons are resolved using THG signal centered at ~433 nm. Collagen is a primary constituent of the peripheral nerve sheath and extracellular matrix encasing individual axons; dysregulation in collagen production impacts nerve regeneration^20,21^. Myelin is a lipid-rich substance that envelops select central and peripheral nervous system axons, providing mechanical protection and electrical insulation for rapid and efficient propagation of action potentials. Dysregulation of myelin production is a hallmark of manifold neurological disease states^22^.

In addition to their use in NLOM, excitation wavelengths around 1300 nm enable three-photon excitation of popular green and yellow fluorescent proteins (GFP, YFP)^23^ and two-photon excitation of several red-fluorescent-dyes^24^. Though use of wavelength-tunable lasers enables excitation of a wide variety of biological markers, use of a single excitation wavelength reduces microscope complexity and cost, while avoiding image distortion arising from chromatic offset inherent with use of multiple excitation wavelengths. Herein, we also demonstrate the utility of this novel fiber-laser-based multiphoton microscope for stain-free multi-color imaging of peripheral nerve from transgenic mice expressing YFP linked to the *thy1* promoter. In this latter application, three-photon-excitation signal with emission maximum at 527 nm delineates the axoplasm of individual axons, while SHG and THG signals delineate neural connective tissue and myelin sheaths, respectively.

## Results

A custom multiphoton microscope (Figs. 1 and 2; see “Methods” section) for multicolor imaging of peripheral nerve was designed and assembled. Here, a custom fiber laser system with a tunable wavelength range from 1045 to 1320 nm was optimized at a single wavelength (~1300 nm), selected to minimize biological sample damage while allowing for deep NLOM imaging. The use of 1300 nm permits simultaneous multiphoton excitation and multiharmonic generation microscopy, while avoiding undesirable two-photon absorption from endogenous fluorophores (namely, NADH and FAD), with subsequent cross-talk and photodamage. System performance and laser parameters including point spread function, pulse energy, and duty cycle were optimized to enhance image contrast and acquisition time while minimizing photodamage. The fs pulse width employed was optimized for multiphoton signal intensity. The PSF of the system was measured using 175 nm blue fluorescent beads; the results for the lateral (0.65 μm) and axial (2.9 μm) directions are shown in Fig. 3. Table 1 in the Methods section summarizes the specifications of the multiphoton microscope. For calibration and PSF measurements, the fiber laser was operated at a repetition rate of 1 MHz. For biological sample imaging, the repetition rate of the laser was increased to 10 MHz to improve scanning efficiency. In both instances the maximum pulse energy at the sample plane was ~10 nJ.

**Figure 1.**
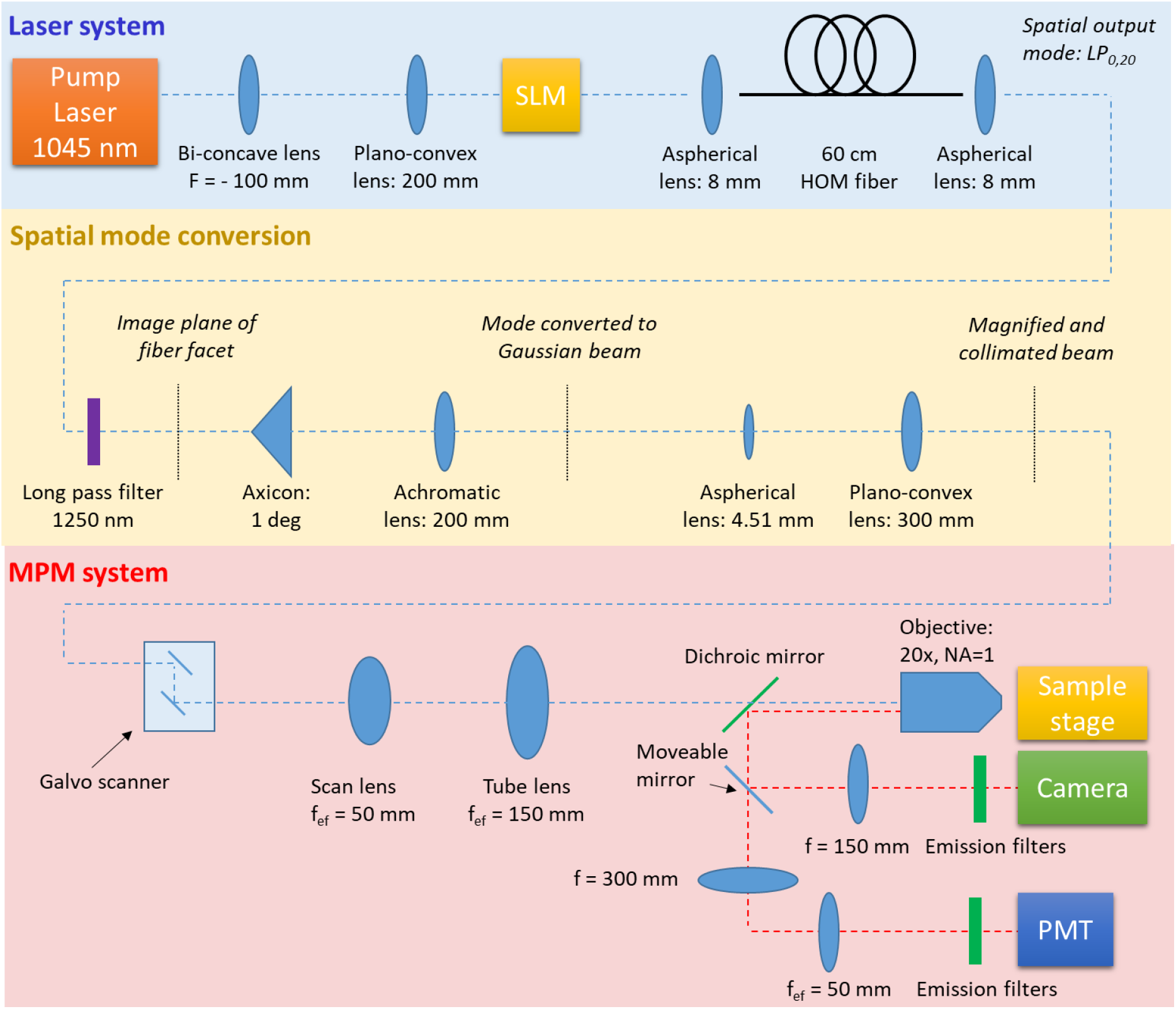
Experimental setup. The hardware employed comprises the laser system (blue shaded region), spatial mode conversion (yellow shaded region), and the MPM system (red shaded region).

**Figure 2.**
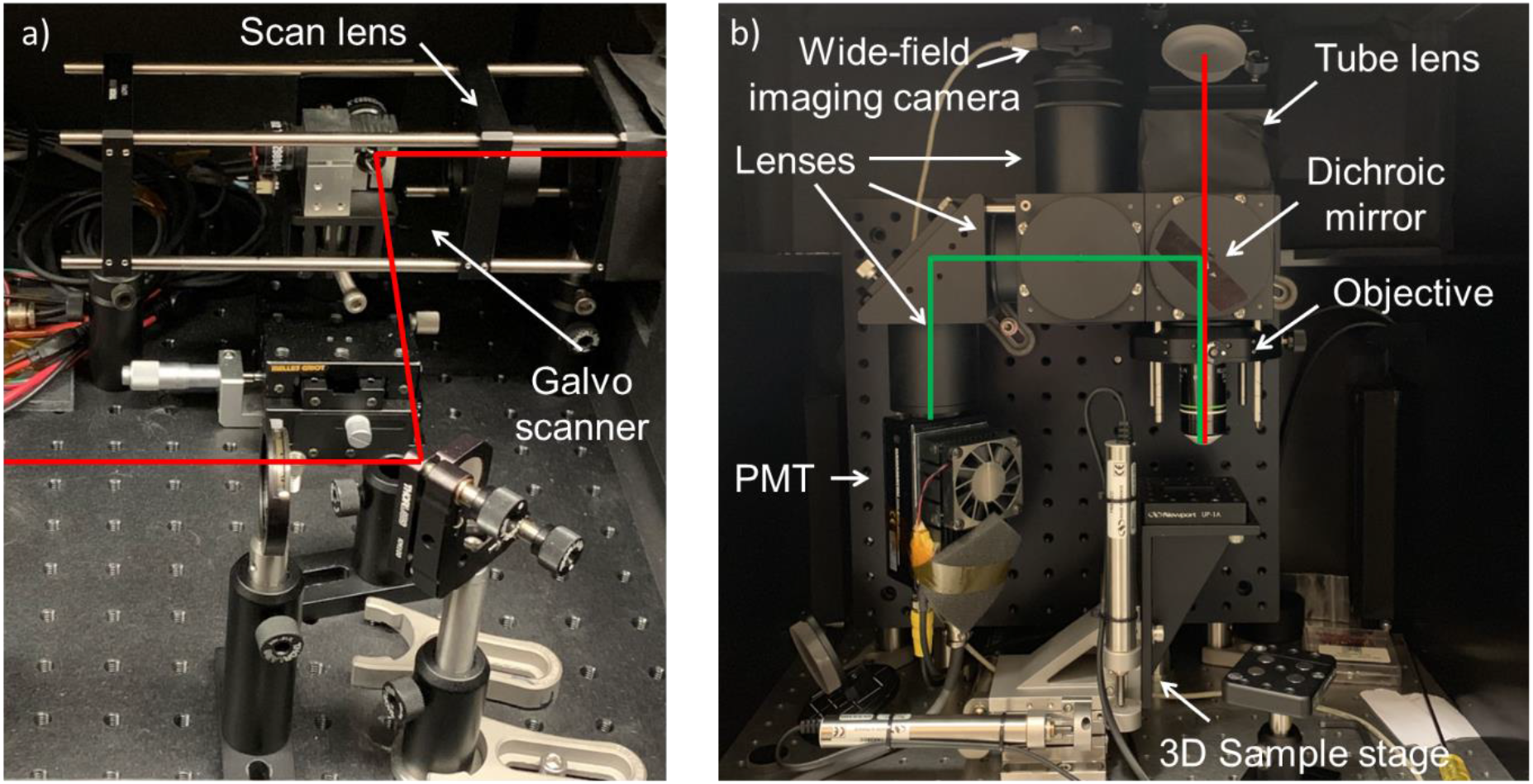
Images of the microscope. Red line indicates path of the excitation beam. Green line indicates collected signal from the sample. A mirror positioned at 45 degrees incidence is located between the scan and tube lens and directs the beam down towards the optical table. (**A**) Image of the raised breadboard with the galvo-scanner and scan lens. (**B**) Side-view image of the microscope.

**Figure 3.**
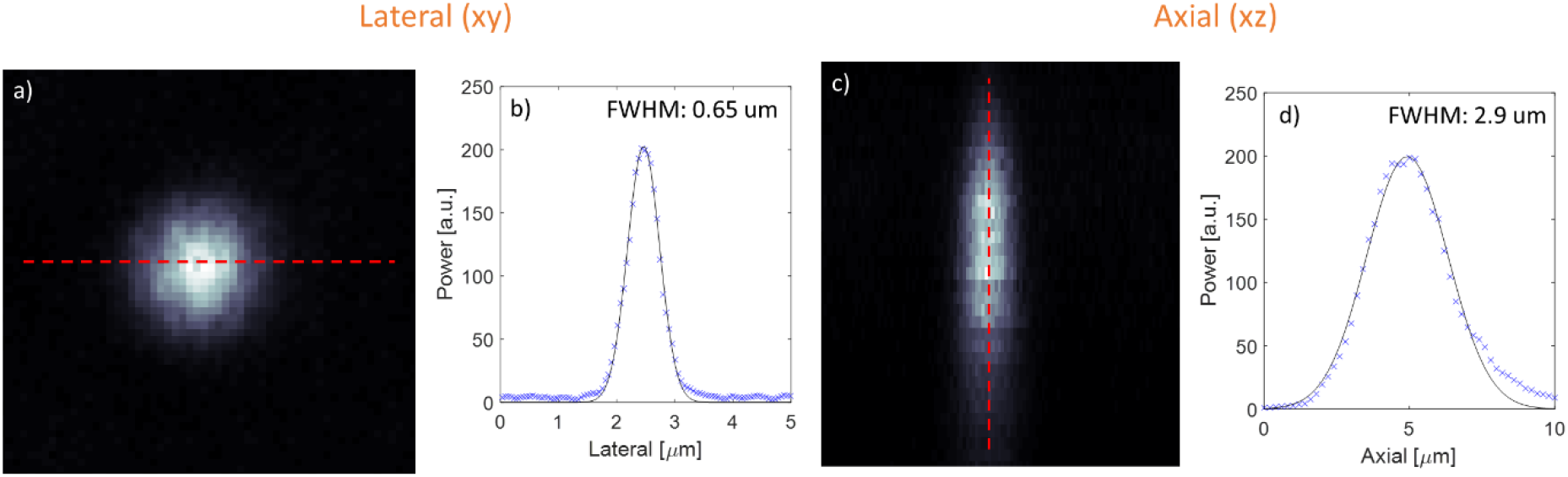
Point spread function measurements. (**A**) The lateral FWHM is 0.65 μm. (**B**) The axial FWHM is 2.9 μm.

**Table 1:** Performance of MPM system.

| Specification | Value |
| --- | --- |
| **Wavelength** | ~1300 nm |
| **Pulse energy (post objective)** | 10 nJ |
| **Pulse duration (post objective)** | 114 fs |
| **Peak power (post objective)** | 87 kW |
| **Average power (post objective)** | 100 mW |
| **Repetition rate** | 10 MHz |
| **MPM system loss** | 6.6 dB |
| **Lateral PSF (FWHM)** | 0.65 $\mu\text{m}$ |
| **Axial PSF (FWHM)** | 2.9 $\mu\text{m}$ |
| **Field-of-view** | 1 x 1 mm |
| **Spatial mode** | Gaussian |

The microscope was used for SHG and THG epidetection imaging of human and murine peripheral nerve. Figure 4A-B shows images of axially frozen-sectioned human obturator nerve obtained sequentially with different band-pass filters (BPF). For this label-free sample, the SHG signal (617/73 nm BPF) was generated from collagen in the three nerve sheath layers (endoneurium, perineurium, epineurium). Collagen is the main tissue component responsible for SHG signal in peripheral nerve tissue. For validation of label-free SHG imaging, picrosirius red was employed on separate tissue sections as an efficient alternative to immunohistochemistry for high-specificity labelling of collagen-dense structures. Picrosirus red-stained collagen was imaged using brightfield microscopy^25^, as shown in Fig. 4C. THG signal (420/50 nm BPF) was predominately generated from the myelin sheaths of myelinated axons (Fig. 4B), demonstrating near identical imaging results compared to widefield fluorescence imaging of myelin-specific dye stained sections, as shown in Fig. 4D. Though THG signal is predominantly emitted in the forward direction, there is sufficient backward-scattering in turbid media (such as peripheral nerve) to obtain sufficient signal using epi-detection as demonstrated herein and by other groups^11,26^. The combination of photobleaching-free SHG and THG images reveal complementary information allowing characterization of nerve morphology in unstained tissues, as shown in Fig. 4E. Figure 4F shows myelinated axon quantification performed through segmentation of a THG image using commercial machine learning software^27^.

**Figure 4.**
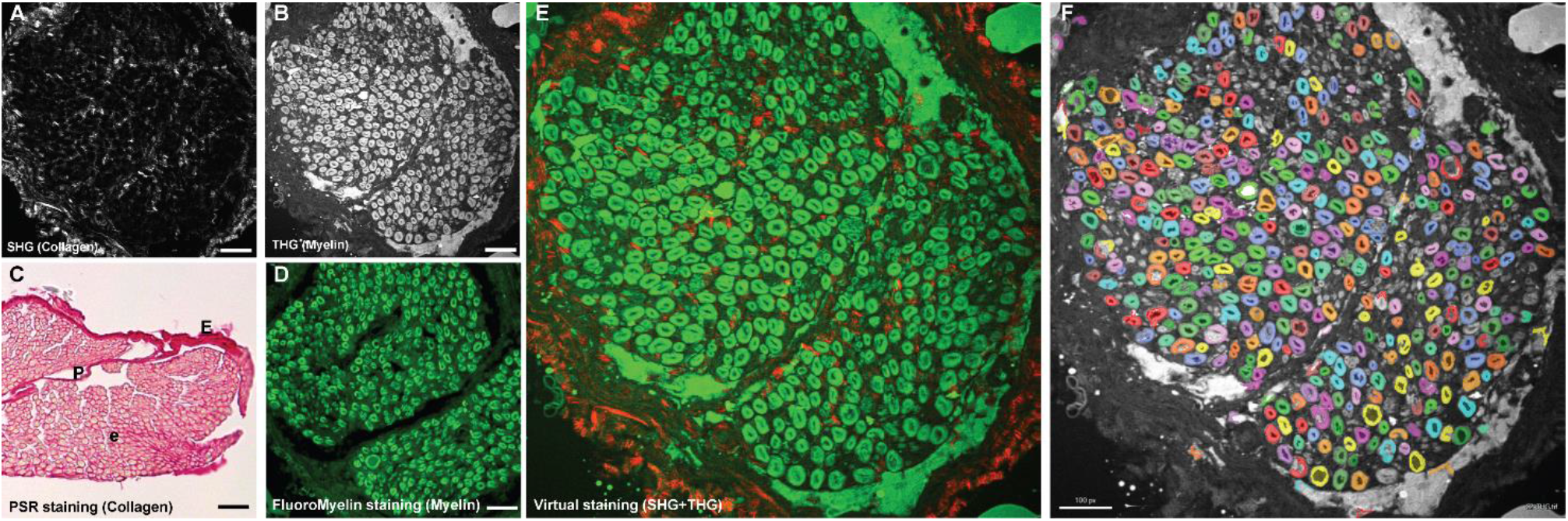
Label-free imaging of ex vivo human obturator nerve cross-sections using a novel 1300 nm fiber laser-powered multiphoton microscope. (**A**) Second-harmonic generation (red) imaging of collagen on neural connective tissue. (**B**) Third-harmonic generation (green) imaging of myelinated fibers. (**C**) Picrosirius red staining of myelinated fibers (conventional light microscopy). (**D**) Myelin-specific dye (FluoroMyelin Green, Molecular Probes, Eugene, OR) staining of cross-section (widefield fluorescence microscopy). (**E**) Merged image combining SHG and THG signals. (**F**) Automated histomorphometry of peripheral nerve human obturator nerve section imaged by THG. Scale bar 60 μm. The FOV is 380 × 380 μm and the scale bar is 50 μm.

Figure 5 demonstrates imaging of a sciatic nerve cross-section from a Thy1-YFP mouse using the multiphoton microscope developed herein. Three-photon-excitation fluorescence images were collected using a third filter (535/60 nm BPF) to separate them from SHG and THG signals. Fig. 5A-C demonstrates THG signal from myelin, SHG signal from collagen, and three-photon fluorescence excitation signal from YFP-labelled axons. In Fig. 5D the three images are merged to obtain a multicolor image.

**Figure 5.**
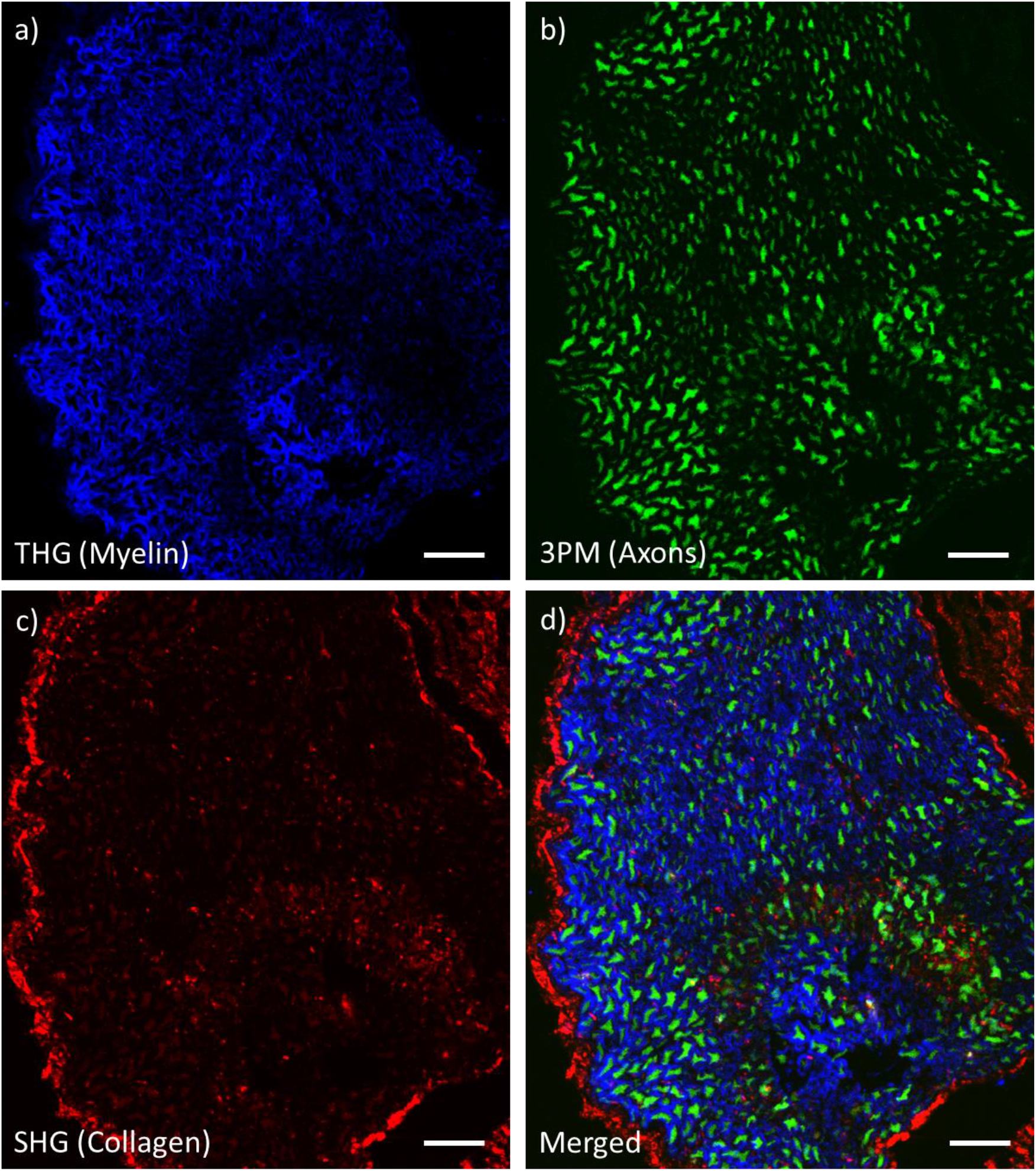
Multiphoton-microscopy image of ex vivo sciatic nerve of a Thy1-YFP mouse. FOV is 210 × 240 μm. Scale bars are 25 μm. (**A**) Third-harmonic generation (blue) shows the myelinated fibers. (**B**) Three-photon excited fluorescence of YFP (green) shows axons (**C**) Second-harmonic generation (red) shows collagen fibers. (**D**) Merged image of THG, 3PE, and SHG signals.

To assess whether this novel 1300 nm fiber-based multiphoton microscope could potentially be employed for in vivo imaging of mammalian nerve, wet mount murine sciatic nerve from a Thy1-YFP mouse was imaged. Figure 6 demonstrates the results for 3PE (Fig. 6A, YFP) and THG imaging (Fig 6B, myelin) using non-descanned epidetection at a depth of 120 μm into the tissue. The two images are merged in Fig. 6C.

**Figure 6.**
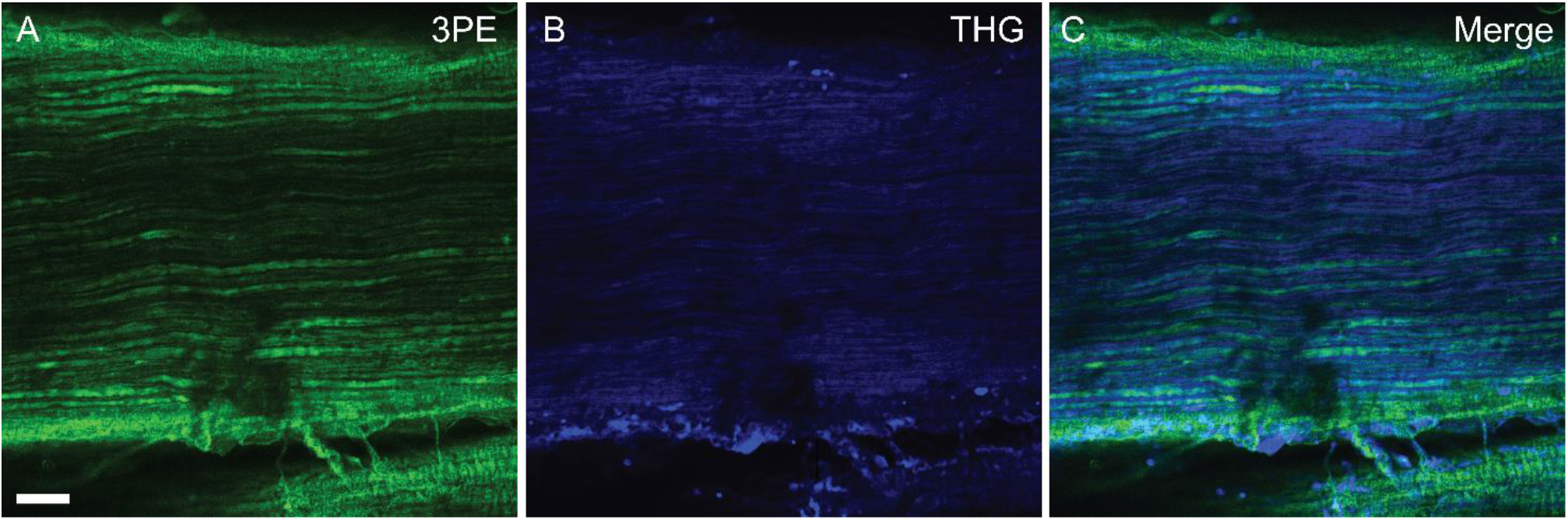
Multiphoton-microscopy images of wet mounted sciatic nerve of a Thy1-YFP mouse at a depth of 120 μm. (**A**) Three-photon excited fluorescence of YFP (green) shows axons (**B**) Third-harmonic generation (blue) signal shows the myelinated fibers. (**C**) Merged image of 3PE and THG signals. 50 frame averages were used to enhance image quality. FOV is 480 × 480 μm. Scale bars is 50 μm.

To estimate the nonlinear order and required power for multiphoton imaging, the dependence of the signal on excitation power was measured. The plot of signal strength versus excitation power is shown in Fig. 7, along with fits. The black-dashed lines represent the power order as a free-fitting parameter, while the red-dashed lines represent fits with fixed power order. The fits show excellent agreement with the data, indicating capture of appropriate emission. Excitation power exhibits a second-order increase in the SHG signal (x1.90) and third-order increase in the 3PE (x2.99) and THG (x3.24) signals.

**Figure 7.**
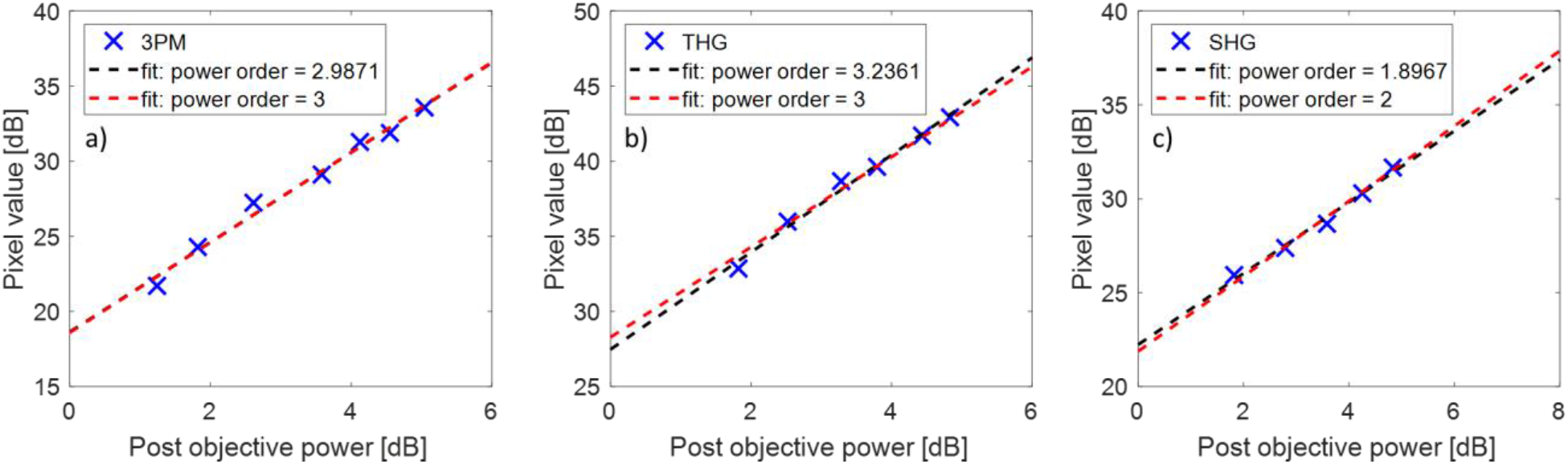
Power dependence of the three signals measured in Fig. 4. (**A**) Three-photon signal. (**B**) THG signal. (**C**) SHG signal.

## Discussion

Herein, we developed a novel microscope based on a single 1300 nm excitation source for multiharmonic and multi-photon excitation microscopy. We demonstrated the utility of the microscope in label-free imaging of peripheral nerve frozen sections of healthy murine sciatic nerve and healthy human obturator nerve, and compared results against diffraction-limited techniques. Three-photon fluorescence, SHG, and THG imaging herein was achieved using a cost-efficient custom fiber-based high-powered near-infrared femtosecond laser source. Despite a long illumination wavelength at 1300 nm, adequate SHG signal to highlight collagen-rich endoneurium, perineurium, and epineurium was obtained via non-descanned epidetection. Additional sections were imaged using a commercial femtosecond laser at 1300 nm (Spectra-Physics Insight X3, Newport Corp, Newport, NJ) on a commercial multiphoton microscope (TrimScope II, LaVision BioTech GmbH). Despite identical PMT and band-pass filters and similar epi-collection non-descanned optical paths, insufficient signal was captured to resolve THG and 3PE signals (Supplemental Figure 1).

The combination of SHG and THG imaging using a novel single fiber femtosecond laser source at 1300 nm represents means for label-free imaging of peripheral nerve with potential implications in diagnosis of diseases impacting myelin regulation and for guiding intraoperative decision making in nerve transfer procedures^9^. In vivo THG imaging of myelin sheaths of peripheral nerve axons carries diagnostic potential and has been previously demonstrated using a 1180 nm laser illumination source^11^. Though THG imaging of brain tissue has been achieved at depths up to a few hundred microns, imaging of peripheral axons in intact peripheral nerve is challenging due to the highly scattering properties of the epineurial sheath. Heretofore, imaging of myelinated axons within intact mammalian peripheral nerve has been limited to 30-50 μm depth. Herein, we achieved an imaging depth exceeding 100 μm in whole nerve using longer wavelength excitation at 1300 nm and a high resolution objective (20X 1.0 NA). Multiphoton microscopy can be implemented on epi and transmission-detection geometries^14^, where for in vivo imaging epidetection is a necessity. In comparison to illumination below 1200 nm, longer illumination wavelengths for THG imaging yield longer wavelength emission in the visible (>400 nm) as opposed to ultraviolet (< 400 nm) domains, lessening undesirable signal absorption by tissue constituents and permitting deeper imaging. Such longer illumination wavelengths also increase the penetration depth of SHG imaging of collagen^28^.

Future work will examine means to optimize lateral and axial resolution in multiphoton microscopy. Epidection NLOM imaging signal is typically collected via non-descanned detection, using a large area detector positioned close to the objective lens to minimize signal loss along the collection path. Alternatively, escanned confocal detection may potentially be employed to minimize background signal and increase axial resolution, though loss of signal along the descanned detection path and pinhole may negate potential gains in the signal-to-background ratio (SBR). In place of a pinhole and single point detector, an array of photodetectors (e.g. Airyscan detection) paired with deconvolution algorithms could be alternatively employed to optimize descanned detection signal by capturing photons that would otherwise be discarded by the pinhole^29^. Spatial resolution could be further enhanced by digital pixel reassignment or blind image reconstruction, as previously reported for 2PE microscopy^30,31^.

## Methods

The experimental setup is shown in Figs. 1 and 2. It consists of three elements, the laser system (blue shaded region), spatial mode conversion (yellow shaded region), and the MPM system (red shaded region), which are described below. For more details on the laser system and spatial mode conversion please see *Rishøj et al*^16^.

### Laser system

The blue shaded region of Fig. 1 is the laser system, wherein a 100 fs pulse at 1045 nm from the pump laser (Y-Fi, KMLabs, Boulder, CO) is converted to ~1300 nm. A telescope magnifies the beam before propagation onto a spatial light modulator (SLM), which encodes a transverse spatial phase onto the incident Gaussian beam to ensure excitation of a single pure mode in the multimode fiber^32^. Here a LP_0,21_ mode is excited in the custom 54 cm higher-order-mode fiber (core diameter of 97 μm and numerical aperture of 0.34). A combination of soliton self-frequency shifting (SSFS) and the newly discovered effect of soliton-self mode conversion (SSMC) leads to output pulses at 1317 nm, with pulse energies of 80 nJ in the LP_0,20_ mode^16,33^. The LP_0,20_ mode resembles a truncated Bessel beam comprising a central peak surrounded by 19 concentric rings. Using an autocorrelator, the pulse duration was measured at 74 fs, corresponding to a peak power of approximately 1.1 MW. Based on the measured spectral bandwidth, it was found that the pulses are nearly transform-limited, an expecting finding considering the pulses are solitons.

### Spatial mode conversion

In the yellow shaded region of the experimental setup shown in Fig. 1, a long-pass filter initially removes the residual pump light. Next, the LP0,20 fiber output mode is converted to a Gaussian-like beam using an axicon and lens, with a measured power conversion efficiency of 81% (theoretically 86%)^16^. The following two lenses magnify and collimate the Gaussian beam to a beam diameter of 5 mm (FWHM). At this point, the pulse duration was measured at 100 fs using an autocorrelator. Though bessel beams may be used to extend the depth of focus and for fast volumetric imaging of sparse samples,^23^ herein a Gaussian beam was utilized as it is more energy-efficient for imaging histological sections of excised nerves.

### Multiphoton microscope

The multiphoton microscope is shown in the red shaded region of Fig. 1. The system was designed de novo around the aforementioned 1300 nm fiber laser, for cost efficiency and performance optimization. Galvo mirrors (GVS002, Thorlabs, Newton, NJ) are used to scan the beam. The scan and tube lenses comprise of several air-spaced commercial lenses from Thorlabs. The optical design was optimized in Zemax to minimize aberrations, resulting in diffraction-limited performance for optical ray angles below 11.3 degrees. The scan lens consists of two LA1541-C lenses and a LA1031-C lens and the tube lens consists of two AC508-300-C. The signals are epi-collected by a 20x (NA=1) objective lens (XLUMPLFLN20XW, Olympus, Tokyo, Japan), and separated by a 775 nm long-pass dichroic mirror (FF775-Di01, Semrock Inc., Rochester, NY), and then detected sequentially using emission filters by a PMT (H7422-40, Hamamatsu Photonics, Hamamatsu, Japan). The collection path was designed in Zemax from commerical lenses (ThorLabs) to ensure maximum collection efficiency. The first lens (ThorLabs LA1256-A) is followed by an air-spaced lens consisting of three plano-convex lenses (ThorLabs LA1417-A). This optical design ensures a full collection field-of-view (FOV) of approximately 1 mm in diameter. The sample stage enabled coarse lateral movement and axial scanning using a 3D stepper motor (TRA25CC, Newport Corp., Irvine, CA). A moveable silver mirror downstream of the dichroic mirror enables optional widefield imaging. The pre-amp is a FEMTO DHPCA-100 (FEMTO Messtechnik GmbH, Berlin, Germany), and the DAQ card is an NI PCIe-6353 (National Instruments Corp., Austin, TX). Several precautions were taken to avoid crosstalk between images. Samples were excited using the same wavelength and pulse duration, but images were sequentially collected beginning with the signal requiring the least pulse energy (i.e., SHG) to minimize photodamage. Images were collected with specific band pass-filters to isolate SHG, THG and 3PE emission signals. The power dependency of the three signals was verified by collecting emission as a function of illumination beam power. The microscope system was controlled using ScanImage and the images were processed using ImageJ software (Fiji Distribution, Version 1.52e)^34,35^. Frame averaging was used to further minimize noise. Colors were arbitrarily assigned to the grayscale images. The pulse duration was measured using an autocorrelator after the objective lens at 114 fs. A summary of the optical performance of the MPM system is provided in Table 1.

### Biological samples

Experimental protocols were approved by the Mass Eye and Ear Internal Review Boards, and all methods were carried out in accordance with relevant guidelines and regulations. Fresh frozen sections of healthy human motor branch of obturator nerve and transgenic mice sciatic nerve were employed. Adult human nerve samples (patient age > 18 years) were obtained fresh at time of free-tissue transfer for smile reanimation after informed consent among patients undergoing facial reanimation procedures. Sciatic nerves from Thy1-YFP mice were harvested in accordance with a Massachusetts Eye and Ear Institutional Animal Care and Use Committee approval. Specifically, animals were placed in an induction chamber and placed under general anesthesia by inhalation of 2% isoflurane in 0.6 L/min O_2_. Once unconscious, isoflurane was increased to 5% and continued until cessation of breathing was noted for one minute, followed by immediate cervical dislocation. Nerves were immediately harvested from the carcasses, and fixed by immersion in 2% phosphate-buffered paraformaldehyde. Sectioned were imaged wet-mount or following sectioned. Sectioned samples underwent overnight cryoprotection in sucrose solution, cryosectioning at 1-2 μm, and staining using a non-toxic myelin specific dye prior to mounting and imaging using a previously described protocol^9^.

## Acknowledgements

The authors would like to thank Anderson Chen (Boston University) for helping with Zemax simulations and Suresh Mohan (Mass Eye and Ear) for sample preparation. A portion of this work was supported by a generous gift from the Berthiaume Family to the Surgical Photonics & Engineering Laboratory at Mass Eye and Ear. The work was also funded by National Institute of Health (1R21EY026410-01), Air Force Office of Scientific Research (FA9550-14-1-0165), and Office of Naval Research (N00014-17-1-2519).

## Author contributions

L.R. and S.R. conceived the idea and designed the experiments. I.C.H. and N.J. prepared the biological samples. L.R. and I.C.H. performed the experiments. L.R., I.C.H, N.J. and S.R. analyzed the data. L.R. and I.C.H wrote the manuscript, and N.J. and S.R. performed critical revisions. All authors read and approved the final manuscript.

## Additional information

### Competing financial interests

The authors declare no competing financial interests.

## Notes

### Competing Interest Statement

The authors have declared no competing interest.

## References

1. Terzis, J. K., Dryer, M. M. & Bodner, B. I. Corneal neurotization: A novel solution to neurotrophic keratopathy. Plast. Reconstr. Surg. (2009) doi:10.1097/PRS.0b013e3181904d3a.

2. Jowett, N. & Pineda, R. Corneal neurotisation by great auricular nerve transfer and scleral-corneal tunnel incisions for neurotrophic keratopathy. British Journal of Ophthalmology (2018) doi:10.1136/bjophthalmol-2018-312563.

3. Elbaz, U., Bains, R., Zuker, R. M., Borschel, G. H. & Ali, A. Restoration of corneal sensation with regional nerve transfers and nerve grafts: A new approach to a difficult problem. JAMA Ophthalmol. (2014) doi:10.1001/jamaophthalmol.2014.2316.

4. O’Brien, B. M., Franklin, J. D. & Morrison, W. A. Cross-facial nerve grafts and microneurovascular free muscle transfer for long established facial palsy. Br. J. Plast. Surg. (1980) doi:10.1016/0007-1226(80)90013-2.

5. Transfer, F. M. & Nerve, F. Functional muscle transfer and the variance of reinnervating axonal load: part I. Peripheral nerves. 1570–1577 doi:10.1097/PRS.0b013e31816fda3e.

6. MacQuillan, A. H. & Grobbelaar, A. O. Functional muscle transfer and the variance of reinnervating axonal load: part II. Peripheral nerves. Plast. Reconstr. Surg. (2008).

7. Terzis, J. K., Wang, W. & Zhao, Y. Effect of axonal load on the functional and aesthetic outcomes of the cross-facial nerve graft procedure for facial reanimation. Plast. Reconstr. Surg. (2009) doi:10.1097/PRS.0b013e3181babb93.

8. Hembd, A. et al. Facial Nerve Axonal Analysis and Anatomical Localization in Donor Nerve: Optimizing Axonal Load for Cross-Facial Nerve Grafting in Facial Reanimation. in Plastic and Reconstructive Surgery (2017). doi:10.1097/PRS.0000000000002897.

9. Wang, W., Kang, S., Coto Hernández, I. & Jowett, N. A Rapid Protocol for Intraoperative Assessment of Peripheral Nerve Myelinated Axon Count and its Application to Cross-Facial Nerve Grafting. Plast. Reconstr. Surg. (2018) doi:10.1097/prs.0000000000005338.

10. Freudiger, C. W. et al. Label-free biomedical imaging with high sensitivity by stimulated Raman scattering microscopy. Science (2008) doi:10.1126/science.1165758.

11. Lim, H. et al. Label-free imaging of Schwann cell myelination by third harmonic generation microscopy. Proc. Natl. Acad. Sci. 111, 18025–18030 (2014).

12. Denk, W., Strickler, J. H. & Webb, W. W. Two-photon laser scanning fluorescence microscopy. Science 248, 73–6 (1990).

13. Diaspro, A., Chirico, G. & Collini, M. Two-photon fluorescence excitation and related techniques in biological microscopy. Quarterly Reviews of Biophysics (2005) doi:10.1017/S0033583505004129.

14. Débarre, D. et al. Imaging lipid bodies in cells and tissues using third-harmonic generation microscopy. Nat. Methods (2006) doi:10.1038/nmeth813.

15. Genthial, R. et al. Label-free imaging of bone multiscale porosity and interfaces using third-harmonic generation microscopy. Sci. Rep. (2017) doi:10.1038/s41598-017-03548-5.

16. Rishøj, L., Tai, B., Kristensen, P. & Ramachandran, S. Soliton self-mode conversion: revisiting Raman scattering of ultrashort pulses. Optica (2019) doi:10.1364/optica.6.000304.

17. Zinter, J. P. & Levene, M. J. Maximizing fluorescence collection efficiency in multiphoton microscopy. Opt. Express (2011) doi:10.1364/oe.19.015348.

18. Yildirim, M., Sugihara, H., So, P. T. C. & Sur, M. Functional imaging of visual cortical layers and subplate in awake mice with optimized three-photon microscopy. Nat. Commun. (2019) doi:10.1038/s41467-018-08179-6.

19. Rishoj, L., Hernandez, I. C., Jowett, N. & Ramachandran, S. Multiharmonic Imaging of Human Peripheral Nerves using a 1300 nm Ultrafast Fiber Laser. in Conference Proceedings - Lasers and Electro-Optics Society Annual Meeting-LEOS (2020).

20. Chen, P., Cescon, M., Megighian, A. & Bonaldo, P. Collagen VI regulates peripheral nerve myelination and function. FASEB J. (2014) doi:10.1096/fj.13-239533.

21. Koopmans, G., Hasse, B. & Sinis, N. Chapter 19 The Role of Collagen in Peripheral Nerve Repair. International Review of Neurobiology (2009) doi:10.1016/S0074-7742(09)87019-0.

22. Duncan, I. D. & Radcliff, A. B. Inherited and acquired disorders of myelin: The underlying myelin pathology. Experimental Neurology (2016) doi:10.1016/j.expneurol.2016.04.002.

23. Chen, B. et al. Rapid volumetric imaging with Bessel-Beam three-photon microscopy. Biomed. Opt. Express (2018) doi:10.1364/boe.9.001992.

24. Wang, T. et al. Three-photon imaging of mouse brain structure and function through the intact skull. Nat. Methods (2018) doi:10.1038/s41592-018-0115-y.

25. Wegner, K. A., Keikhosravi, A., Eliceiri, K. W. & Vezina, C. M. Fluorescence of Picrosirius Red Multiplexed With Immunohistochemistry for the Quantitative Assessment of Collagen in Tissue Sections. J. Histochem. Cytochem. (2017) doi:10.1369/0022155417718541.

26. Débarre, D., Olivier, N. & Beaurepaire, E. Signal epidetection in third-harmonic generation microscopy of turbid media. Opt. Express (2007) doi:10.1364/oe.15.008913.

27. Coto Hernández, I., Yang, W., Mohan, S. & Jowett, N. Label-free histomorphometry of peripheral nerve by stimulated Raman spectroscopy. Muscle and Nerve (2020) doi:10.1002/mus.26895.

28. Balu, M. et al. Effect of excitation wavelength on penetration depth in nonlinear optical microscopy of turbid media. J. Biomed. Opt. (2009) doi:10.1117/1.3081544.

29. Huff, J., Kleppe, I., Naumann, A. & Nitschke, R. Airyscan detection in multiphoton microscopy: super-resolution and improved signal-to-noise ratio beyond the confocal depth limit. Nat. Methods 3–5 (2018).

30. Sheppard, C. J. R., Castello, M., Tortarolo, G., Vicidomini, G. & Diaspro, A. Image formation in image scanning microscopy, including the case of two-photon excitation. J. Opt. Soc. Am. A (2017) doi:10.1364/josaa.34.001339.

31. Koho, S. V. et al. Two-photon image-scanning microscopy with SPAD array and blind image reconstruction. Biomed. Opt. Express (2020) doi:10.1364/boe.374398.

32. Demas, J., Rishøj, L. & Ramachandran, S. Free-space beam shaping for precise control and conversion of modes in optical fiber. 23, 28531–28545 (2015).

33. Rishøj, L., Tai, B., Kristensen, P. & Ramachandran, S. Characterization of intermodal group index matched soliton interactions leading to mw peak powers at 1300 nm. in 2017 Conference on Lasers and Electro-Optics, CLEO 2017 - Proceedings (2017). doi:10.1364/CLEO_AT.2017.STh3K.2.

34. Schindelin, J. et al. Fiji: An open-source platform for biological-image analysis. Nature Methods vol. 9 676–682 (2012).

35. Schneider, C. A., Rasband, W. S. & Eliceiri, K. W. NIH Image to ImageJ: 25 years of image analysis. Nature Methods (2012) doi:10.1038/nmeth.2089.

