## supplementary for "Multiphoton microscopy for label-free multicolor imaging of peripheral nerve"

Supplemental Fig. 1 demonstrates the lack of signal obtained using a commercial femtosecond laser at 1300 nm owing to insufficient peak power.

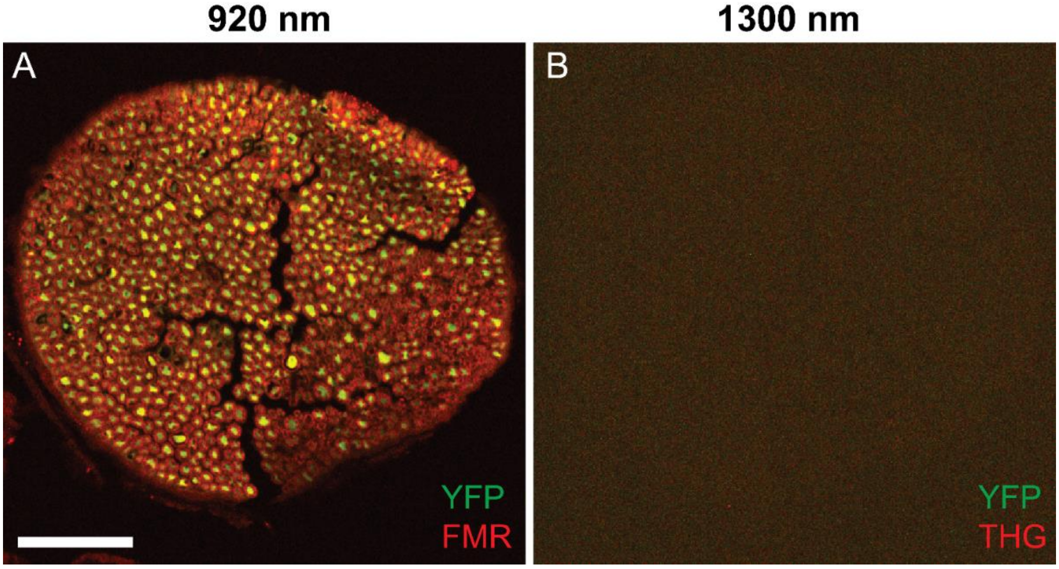


**Suppl. Fig. 1.** Cross-section of the buccal branch of the facial nerve in Thy1-YFP-16 mouse stained with a myelin-specific dye (FluoroMyelin Red®, F34652 Invitrogen, Carlsbad, CA) and imaged using a commercial multiphoton microscope (TrimScope II, LaVision BioTech GmBH) powered by a commercial dual-output femtosecond laser (Insight X3, Spectra Physics). (**A**) Two-photon fluorescence microscopy image (excitation 920 nm) demonstrates axon (YFP) and myelin signal (FMR). (**B**) Same volume imaged at near-maximal output power at 1300 nm demonstrated absent THG signal. Scale bar 100 μm.

**Direct comparison between confocal and NLOM harmonic generation**

A simulation was done for theoretical comparison of diffraction-limited and NLOM harmonic generation techniques (see **Suppl. Fig. 2.**). Herein, theoretical comparison was made between THG and SHG signals at 3x illumination wavelength (e.g. 1300 nm) versus diffraction-limited techniques at 1x illumination wavelength (e.g. 433 nm). Though use of threefold longer illumination wavelengths implies a threefold decrease in resolution, theory predicts slightly lower decrease in resolution owing to the nonlinearity of harmonic imaging.

Supplemental Fig. 2 shows the normalized PSF for confocal microscopy, SHG and THG. Herein the case of no Stokes shift was considered for confocal, and illumination wavelengths for SHG and THG were 2 and 3 times larger the wavelength used for confocal, respectively. The curves are plotted against the cylindrical optical coordinate $v = (2\pi/\lambda) rnsin\alpha$, where n sin α is the numerical aperture and r is the cylindrical radius. Herein was used the theory of multiphoton microscopy described by Sheppard et al[1].

$$I_{conf}(v) = \left[\frac{2J_1(v)}{v}\right]^2 * \left[\frac{2J_1(v)}{v}\right]^2$$

$$I_{SHG}(v) = \left[\frac{2J_1(v/2)}{v/3}\right]^2 * \left[\frac{2J_1(v/2)}{v/3}\right]^2$$

$$I_{THG}(v) = \left[\frac{2J_1(v/3)}{v/3}\right]^2 * \left[\frac{2J_1(v/3)}{v/3}\right]^2 * \left[\frac{2J_1(v/3)}{v/3}\right]^2$$

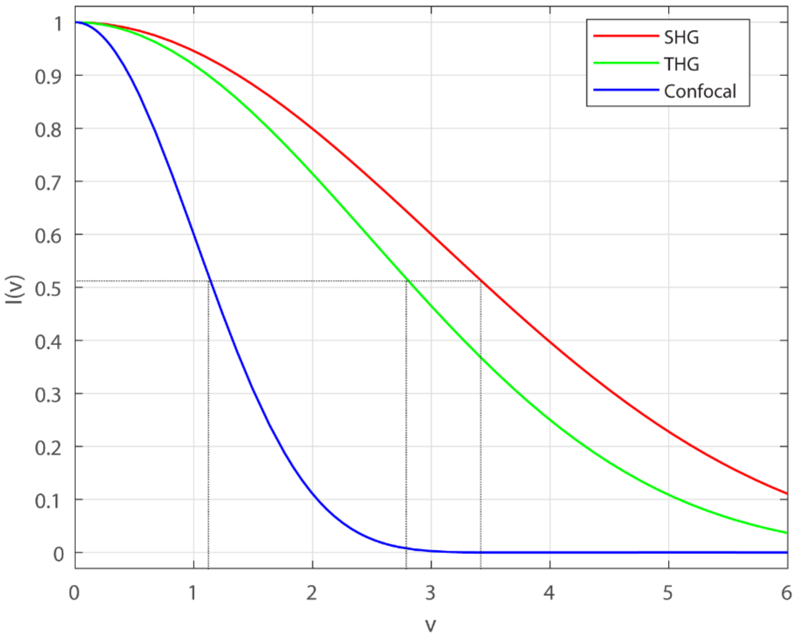


**Suppl. Fig. 2.** Resolution comparison between second and third harmonic generation with confocal microscopy. Theoretical intensity plot as a function of normalized radial optical coordinate (v) of SHG, THG, and single-photon excitation confocal microscopy signals. The resolution of THG and SHG is lower than diffraction-limited techniques by a factor of 2.44 and 2.97, respectively, assuming a three-fold longer illumination wavelength is used for THG and SHG as compared to diffraction-limited imaging.

1. SHEPPARD, C. & GU, M. Image formation in two-photon fluorescence microscopy. *Opt.* (1990).
